# PlantTribes2: tools for comparative gene family analysis in plant genomics

**DOI:** 10.1101/2022.11.17.516924

**Authors:** Eric K. Wafula, Huiting Zhang, Gregory Von Kuster, James H. Leebens-Mack, Loren A. Honaas, Claude W. dePamphilis

## Abstract

Plant genome-scale resources are being generated at an increasing rate as sequencing technologies continue to improve and raw data costs continue to fall; however, the cost of downstream analyses remains large. This has resulted in a considerable range of genome assembly and annotation qualities across plant genomes due to their varying sizes, complexity, and the technology used for the assembly and annotation. To effectively work across genomes, researchers increasingly rely on comparative genomic approaches that integrate across plant community resources and data types. Such efforts have aided the genome annotation process and yielded novel insights into the evolutionary history of genomes and gene families, including complex non-model organisms. The essential tools to achieve these insights rely on gene family analysis at a genome-scale, but they are not well integrated for rapid analysis of new data, and the learning curve can be steep. Here we present PlantTribes2, a scalable, easily accessible, highly customizable, and broadly applicable gene family analysis framework with multiple entry points including user provided data. It uses objective classifications of annotated protein sequences from existing, high-quality plant genomes for comparative and evolutionary studies. PlantTribes2 can improve transcript models and then sort them, either genome-scale annotations or individual gene coding sequences, into pre-computed orthologous gene family clusters with rich functional annotation information. Then, for gene families of interest, PlantTribes2 performs downstream analyses and customizable visualizations including, (1) multiple sequence alignment, (2) gene family phylogeny, (3) estimation of synonymous and non-synonymous substitution rates among homologous sequences, and (4) inference of large-scale duplication events. We give examples of PlantTribes2 applications in functional genomic studies of economically important plant families, namely transcriptomics in the weedy Orobanchaceae and a core orthogroup analysis (CROG) in Rosaceae. PlantTribes2 is freely available for use within the main public Galaxy instance and can be downloaded from GitHub or Bioconda. Importantly, PlantTribes2 can be readily adapted for use with genomic and transcriptomic data from any kind of organism.

## 1. Introduction

A rapid and continuing decline in sequencing costs over the last 30 years has contributed to the generation of massive amounts of transcriptome and genome data for non-model plant species (Barrett et al., 2013; Matasci et al., 2014; Sayers et al., 2018; One Thousand Plant Transcriptomes Initiative, 2019; Marks et al., 2021). Integrating new genomic data from diverse plant lineages in phylogenetic studies can provide the evolutionary context necessary understanding the evolution of gene function (Williams et al., 2014; Yang et al., 2015b; Zhang et al., 2015; Carvalho et al., 2018; Pabón-Mora et al., 2014; Mi et al., 2020; Nagy et al., 2020; One Thousand Plant Transcriptomes Initiative, 2019), resolving species relationships (Timme et al., 2012; Rothfels et al., 2013; Wickett et al., 2014; Zeng et al., 2014; Huang et al., 2016; One Thousand Plant Transcriptomes Initiative, 2019; Hodel et al., 2022; Xiang et al., 2017; Yang et al., 2015a), for accurate identification of orthologous and paralogous genes among species (Sonnhammer and Koonin, 2002; Emms and Kelly, 2019; Derelle et al., 2020; Gabaldón, 2008; Fuentes et al., 2021; Schreiber et al., 2014), and unraveling gene and genome duplications (Jiao et al., 2011, 2012; The Amborella Genome Project, 2013; Li et al., 2015; Bowers et al., 2003; Zwaenepoel and Peer, 2019; Ren et al., 2018; Viruel et al., 2019). However, comparative genomic and phylogenomic analyses typically requires a level of bioinformatic expertise and a scale of computational resources that are inaccessible to many researchers. For instance, a large-scale phylogenomic study may require objective circumscription of representative protein sequences into gene families using a carefully selected set of most appropriate reference genomes. This requires knowledge and skill to assess the quality of available genomic resources as well as an evolutionary perspective to avoid pitfalls that lead to distorted conclusions, such as using a biased selection of reference species or outgroups. In addition, to execute these analytical pipelines, command line skills and the expertise to navigate through and properly set parameters, select appropriate algorithms, and solve potential computation environment conflicts are needed. Although some software (Chen et al., 2020; Tello-Ruiz et al., 2020; Valentin et al., 2020; Bel et al., 2021; Oliveira et al., 2021; Emms and Kelly, 2022) are more user-friendly (*i*.*e*., incorporate a graphical user interface, containerized tools, *etc*.) and have pre-defined parameters suitable for plant research, most others still require custom optimization or are mainly applied to species with small genomes (*i*.*e*., prokaryotes), or non-plant systems (Dunn et al., 2013; Blom et al., 2016; Lanza et al., 2016; Altenhoff et al., 2019; Ebmeyer et al., 2021; Perrin and Rocha, 2021; Pucker et al., 2020; Pucker 2022).

With the goal to improve data accessibility, databases have been created to host curated plant-specific genomic information at different scales, ranging from those including sequenced genomes from diverse plant species (*i*.*e*., PLAZA 5.0, Bel et al., 2021 and Gramene, Tello-Ruiz et al., 2020) to ones focusing on specific plant groups, such as the Genome Database for Rosaceae (GDR, Jung et al., 2018). Major plant databases are reviewed and described by various authors (Martinez, 2016; Chen et al., 2006; Lyons and Freeling, 2008; Wall et al., 2008; Goodstein et al., 2012; Schreiber et al., 2014; Huerta-Cepas et al., 2016; Kriventseva et al., 2018; Mi et al., 2020; Tello-Ruiz et al., 2020; Bel et al., 2021). Some databases also provide gene homology information and computational tools for comparative genomic analysis (Martinez, 2016). However, analysis tools implemented in such databases are typically limited, static, and can only be used to analyze existing data (Tomcal et al., 2013; Sundell et al., 2015; Spannagl et al., 2016; Nakaya et al., 2017; Tello-Ruiz et al., 2020). Some more recent databases contain flexible tools (*i*.*e*., users can select different algorithms), but these are often not scalable (*i*.*e*., many have limitations on data size and number of input sequences). For example, the PLAZA 5.0 database contains 134 carefully selected high-quality plant genomes and provides gene family circumscriptions with rich gene homology and annotation information (Bel et al., 2021). However, users can only upload up to 300 new sequences for the BLAST based gene family search function, and add a maximum of 50 external sequences while running a gene family phylogeny on their webserver (https://bioinformatics.psb.ugent.be/plaza/). These limitations on data input make it infeasible to use these databases to perform genome-scale analyses on new datasets brought by the user.

Other new developments aiming to make complicated bioinformatic analyses accessible to more users are workflow management systems which integrate analytic pipelines and complementary software into readily executable packages, such as SnakeMake (Mölder et al., 2021), Nextflow (Tommaso et al., 2017), Pegasus (Deelman et al., 2015), Galaxy Workbench (Afgan et al., 2018), and others. Of those, the Galaxy Workbench is an open-source web-based software framework that aims to make command-line tools accessible to users without informatic expertise (Afgan et al., 2018), and is popular among biologists. Galaxy implements several comparative genomic tools developed by the bioinformatics community (Darling et al., 2010; Thanki et al., 2018).

Such a web-based framework provides a simplified way to execute standardized analyses and workflows. They can also eliminate the complex administrative and programming tasks inherent in performing big data analyses via batch processing on the command line, and greatly simplify record keeping and re-implementation of complex analysis processes. Often, scientists can perform analyses with either existing or user implemented tools from a web browser. Additionally, individual institutions can link these web-based platforms to their own high-performance computing resources, allowing computationally intensive analysis not always possible on a purely web-based platform.

In an effort to address these accessibility and computational challenges in genome-scale research and to take advantage of the Galaxy environment, we developed PlantTribes2, a gene family analysis framework that uses objective classifications of annotated protein sequences from genomes or transcriptomes for comparative and evolutionary analyses of gene families from any type of organism, including fungi, microbes, animals, and plants. An initial version of PlantTribes was developed by Wall et al. (2008), but has become outdated due to several of the previously mentioned limitations. In PlantTribes2, we have completely revamped PlantTribes from a static relational database to a flexible analytical pipeline with all new code, new features, and extensive testing. We have developed a well-documented analytic framework complete with training materials including tutorials and sample datasets. Finally, we worked with the Galaxy community to develop Galaxy wrappers for all of the PlantTribes2 tools (Blankenberg et al., 2014. Supplemental table 1), so they are available on the public server at usegalaxy.org, and can be installed into any Galaxy instance. Finally, we demonstrate genome-scale evolutionary analysis of gene families using PlantTribes2, starting with *de novo* assembled transcriptomes and gene models from whole genome data. Although our examples, sample datasets, and gene family scaffolds are for plants, the pipeline is system agnostic and can be readily used with genome-scale information from any set of related organisms.

## 2. Pipeline implementation

The PlantTribes2 toolkit is a collection of self-contained modular analysis pipelines that use objective classifications of annotated protein sequences from sequenced genomes for comparative and evolutionary analyses of genome-scale gene families. At the core of PlantTribes2 analyses are the gene family scaffolds, which are clusters of orthologous and paralogous sequences from specified sets of inferred protein sequences. The tools interact with these scaffolds, as described below, to deliver the following outputs: (1) predicted coding sequences and their corresponding translations, (2) a table of pairwise synonymous/non-synonymous substitution rates for either orthologous or paralogous transcript pairs, (3) results of significant duplication components in the distribution of synonymous substitutions rates (Ks), (4) a summary table for transcripts classified into orthologous gene family clusters with their corresponding functional annotations, (5) gene family amino acid and nucleotide fasta sequences, (6) multiple sequence alignments, and (7) inferred maximum likelihood phylogenies (Figure 1)

**Figure 1:**
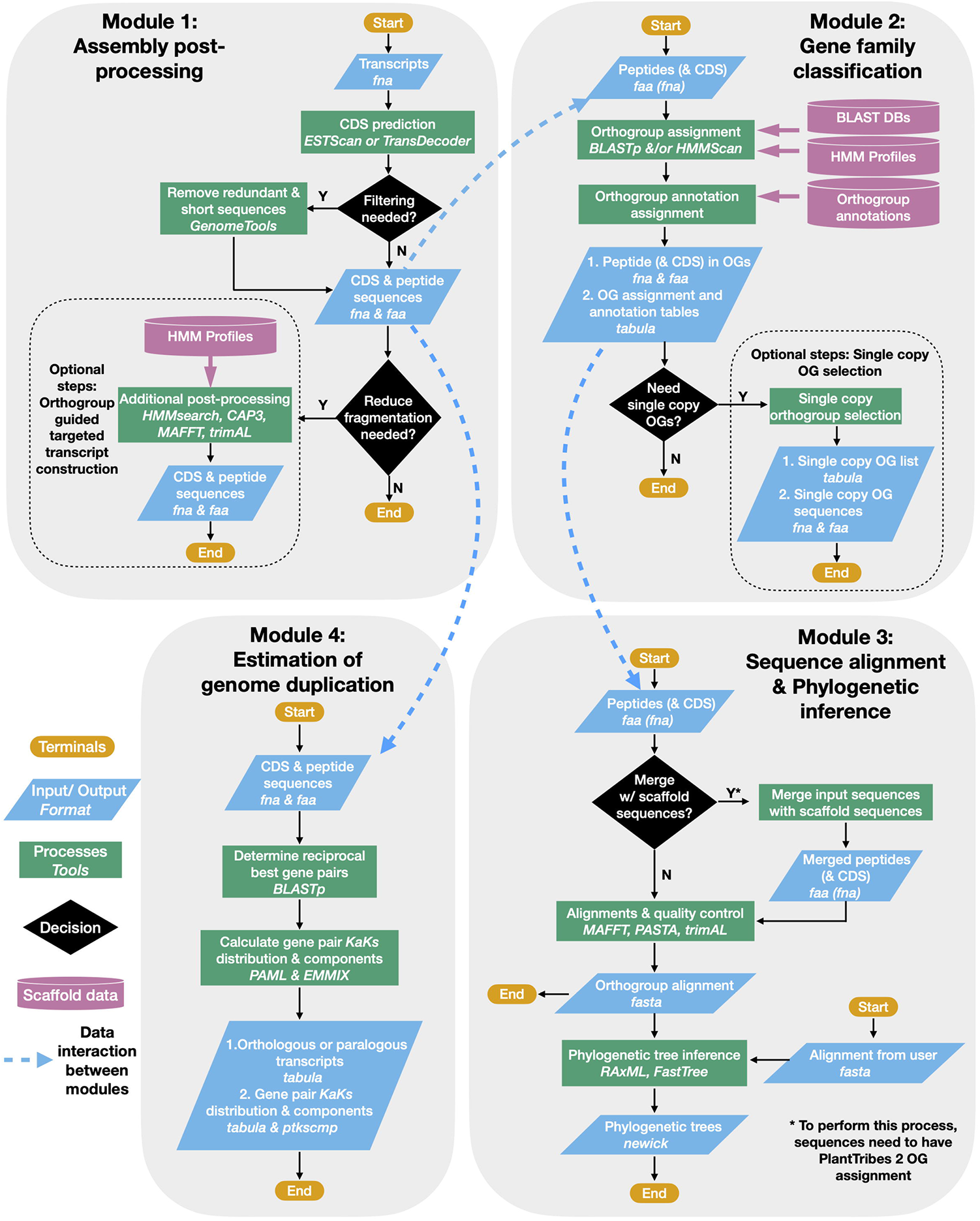
PlantTribes2 analysis workflow. A schematic diagram illustrating the PlantTribes2 modular analysis workflow. (1) A user provides transcripts for post-processing, resulting in a non-redundant set of predicted coding sequences and their corresponding translations (Module 1). (2) The post-processed transcripts (or user provided sequences) are searched against a gene family scaffold blast and/or hmm database(s), and transcripts are assigned into their putative orthogroups with corresponding metadata (Module 2). (3) Classified transcripts are integrated with their corresponding scaffold gene models to estimate orthogroup multiple sequence alignments and corresponding phylogenetic trees (Module 3). Similarly, sequence alignments and phylogeny can be constructed from user provided data. (4) Synonymous substitution rate (Ks) and nonsynonymous substitution rate (Ka) of paralogs from either the post-processed assembly or inferred from the phylogenetic trees are estimated. The Ks results are used to detect large-scale duplication events and in many other evolutionary hypotheses (Module 4).

### 2.1 Gene family scaffolds

The current release of PlantTribes2 (v1.0.4) provides several plant gene family scaffolds (Supplemental table 2) used in previously published and ongoing phylogenomic studies (The Amborella Genome Project et al., 2013; Wickett et al., 2014; Yang et al., 2015b, 2019; Li et al., 2018; Shahid et al., 2018; and Zhang et al., 2022, the companion paper in this issue), including one Monocot focused scaffold (12Gv1.0) and four iterations of generic Angiosperm focused scaffolds (22Gv1.1, 26Gv1.0, 26Gv2.0, and 37Gv1.0). Complete sets of inferred protein-coding genes from plant genomes represented in each of the PlantTribes2 scaffolds were clustered into gene families (*i*.*e*., orthogroups) using at least one of the following protein clustering methods: GFam (clusters of consensus domain architecture) (Sasidharan et al., 2012), OrthoMCL (narrowly defined clusters) (Li et al., 2003; Chen et al., 2006), or OrthoFinder (more broadly defined clusters) (Emms and Kelly, 2015, 2019). Additional clustering of primary gene families was performed using the MCL algorithm (Enright et al., 2002) at 10 stringencies with inflation values from 1.2 to 5.0 to connect distantly, but potentially related orthogroups into larger hierarchical gene families (*i*.*e*., super-orthogroups), as described in Wall et al. (2008). We then annotated each orthogroup with gene function information from biological databases, including Gene Ontology (GO) (Ashburner et al., 2000; Carbon et al., 2019), InterPro/Pfam protein domains (Jones et al., 2014; Blum et al., 2020; Mistry et al., 2020), The Arabidopsis Information Resource (TAIR) (Berardini et al., 2015), UniProtKB/TrEMBL (The UniProt Consortium, 2021), and UniProtKB/Swiss-Prot (The UniProt Consortium, 2021). The final PlantTribes2 scaffold data sets include (1) orthogroups protein coding sequence fasta, (2) orthogroups protein multiple sequence alignments, (3) orthogroups protein HMM profiles, (4) a scaffold protein BLAST database, (5) a scaffold protein HMM profiles database, and (6) templates for analysis pipelines with scaffold metadata.

For custom applications with any focal group of organisms, a detailed description is available on the GitHub repository (https://github.com/dePamphilis/PlantTribes) for how to build a customized PlantTribes2 gene family scaffold. Building custom gene family scaffolds in PlantTribes2 begins with providing unclassified genome-scale gene sets or converting an existing gene family circumscription and corresponding metadata to a format that is compatible with the PlantTribes2 tools. If running on the command line, such externally circumscribed scaffolds can be directly integrated into PlantTribes2 for user-specific gene family analyses. If running on Galaxy, Galaxy administration tools (Blankenberg et al., 2014, Supplemental table 1) are available for installing and maintaining these external scaffolds within a Galaxy instance that provides the PlantTribes2 tools.

### 2.2 Illustrated examples of PlantTribes2 Tools

Here we describe the use of each PlantTribes2 tool and provide examples of outputs using a test dataset containing transcripts from two plant species (details can be found in Supplemental table 3). Detailed step-by-step tutorials using the test data to perform analyses are available for both the Galaxy and the command-line versions of the pipeline.

#### 2.2.1 Assembly post-processing

The *AssemblyPostProcessor* tool is an entry point of a PlantTribes2 analysis when the input data is *de novo* transcripts or gene models in some poorly annotated genomes where predicted coding sequences and corresponding peptides do not match. The *AssemblyPostProcessor* pipeline uses either ESTScan (Iseli et al., 1999) or TransDecoder (Haas et al., 2013) to transform transcripts into putative CDSs and their corresponding amino acid translations. Optionally, the resulting predicted coding regions can be filtered to remove duplicated and exact subsequences using GenomeTools (Gremme et al., 2013). The pipeline is implemented with an additional assembly post-processing method that uses scaffold orthogroups to reduce fragmentation in a *de novo* assembly. Homology searches of post-processed transcripts against HMM-profiles (Eddy, 2011) of targeted orthogroups are conducted using HMMER hmmsearch (Eddy, 2011). After assignment of transcripts to targeted orthogroups, orthogroup-specific gene assembly of overlapping primary contigs is performed using CAP3 (Huang and Madan, 1999), an overlap-layout-consensus assembler. Finally, protein multiple sequence alignments of orthogroups are estimated and trimmed using MAFFT (Katoh and Standley, 2013) and trimAL (Capella-Gutiérrez et al., 2009) respectively, to aid in identifying targeted assembled transcripts that are orthologous to the scaffold reference gene models based on the global sequence alignment coverage. A list of *AssemblyPostProcessor* use cases include: (1) processing *de novo* transcriptome assemblies to improve transcript qualities for downstream analyses (Honaas et al., 2016; Yang et al., 2019; Whittle et al., 2021; and example in section 3.2.1); (2) generating matching coding sequences (CDSs) and peptide sequences in genomes with only mRNA sequences (*i*.*e*., the *Malus domestica* GDDH13 annotation provided only mRNA sequences but not CDSs, Daccord et al., 2017) and gene information gathered from databases lacking a uniformed naming system and processing protocols -for instance, the numbers of CDSs and peptides do not match in the *Pyrus pyrifolia* ‘Cuiguan’ genome, and the peptides are named differently from the CDSs (Gao et al., 2021). The *AssemblyPostProcessor*-generated matching CDS and peptide sequences from the aforementioned *Malus* and *Pyrus* genomes among others provided a good starting point for the comparative genomic analyses described in section 3.2.2 and 3.2.3.

#### 2.2.2 Gene family classification

The *GeneFamilyClassifier* tool classifies gene coding sequences either produced by the *AssemblyPostProcessor* tool or from an external source using BLASTp (Camacho et al., 2009) and HMMER (Eddy, 2011) hmmscan (or both classifiers) into pre-computed orthologous gene family clusters (orthogroups) of a PlantTribes2 scaffold. Classified sequences are then assigned with the corresponding orthogroups’ metadata, which includes gene counts of scaffold taxa, superclusters (super orthogroups) at multiple clustering stringencies, and rich orthogroup annotations from functional genomic databases (as described in section 2.1). Additionally, sequences belonging to single or low-copy gene families that are commonly used in species tree inference can be determined with a built-in command for this tool. Next, the classified input gene coding sequences can be integrated into their corresponding orthogroup’s scaffold gene model files using the *GeneFamilyIntegrator* tool for downstream analyses.

#### 2.2.3 Gene family alignment estimation

The *GeneFamilyAligner* tool estimates protein and codon multiple sequence alignments of integrated orthologous gene family fasta files produced by the *GeneFamilyIntegrator* tool or from an external source. Orthogroup alignments are estimated using either MAFFT’s L-INS-i algorithm (Katoh and Standley, 2013) or the divide and conquer approach implemented in the PASTA (Mirarab et al., 2015) pipeline for large alignments. Optional post-alignment processing includes trimming out sites that are predominantly gaps (Capella-Gutiérrez et al., 2009), removing sequences with very low global orthogroup alignment coverage, and performing realignment of orthogroup sequences following site trimming and sequence removal. In the Galaxy framework, the MSAViewer (Yachdav et al., 2016) plugin allows orthogroup fasta multiple sequence alignments produced by the *GeneFamilyAligner* to be visualized and edited using the Jalview Java Web Start (Waterhouse et al., 2009) (Figure 2).

**Figure 2:**
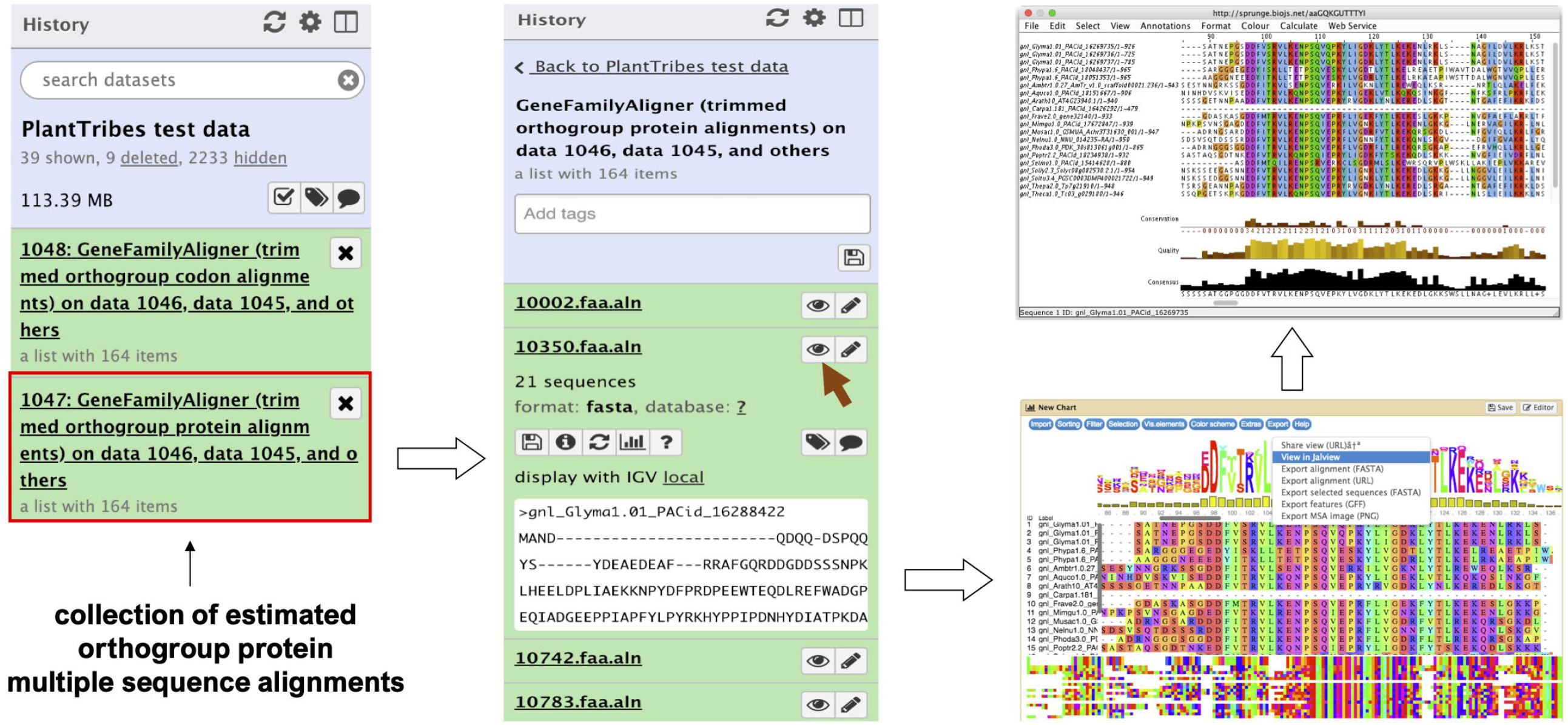
An illustration of an orthogroup multiple sequence alignment produced by the Galaxy PlantTribes2 GeneFamilyAligner tool using the test dataset. Results can be visualized in Galaxy with the MSAViewer visualization plugin and manually edited with Jalview Java Web Start.

#### 2.2.4 Gene family phylogenetic inference

The *GeneFamilyPhylogenyBuilder* tool performs a gene family phylogenetic inference of multiple sequence alignments produced by the *GeneFamilyAligner* tool or from an external source. PlantTribes2 estimates maximum likelihood (ML) phylogenetic trees using either RAxML (Stamatakis, 2014) or FastTree (Price et al., 2010) algorithms. Optional tree optimization includes setting the number of bootstrap replicates for RAxML to conduct a rapid bootstrap analysis, searching for the best-scoring ML tree, and rooting the inferred phylogenetic tree with the most distant taxon in the orthogroup or specified taxa. In the Galaxy framework, either the Phylocanvas plugin (https://phylocanvas.org/) or the PHYLOViZ 2.0 (Nascimento et al., 2016) plugin provides several options for visualizing and rendering the phylogenetic trees produced by the *GeneFamilyPhylogenyBuilder* (Figure 3).

**Figure 3:**
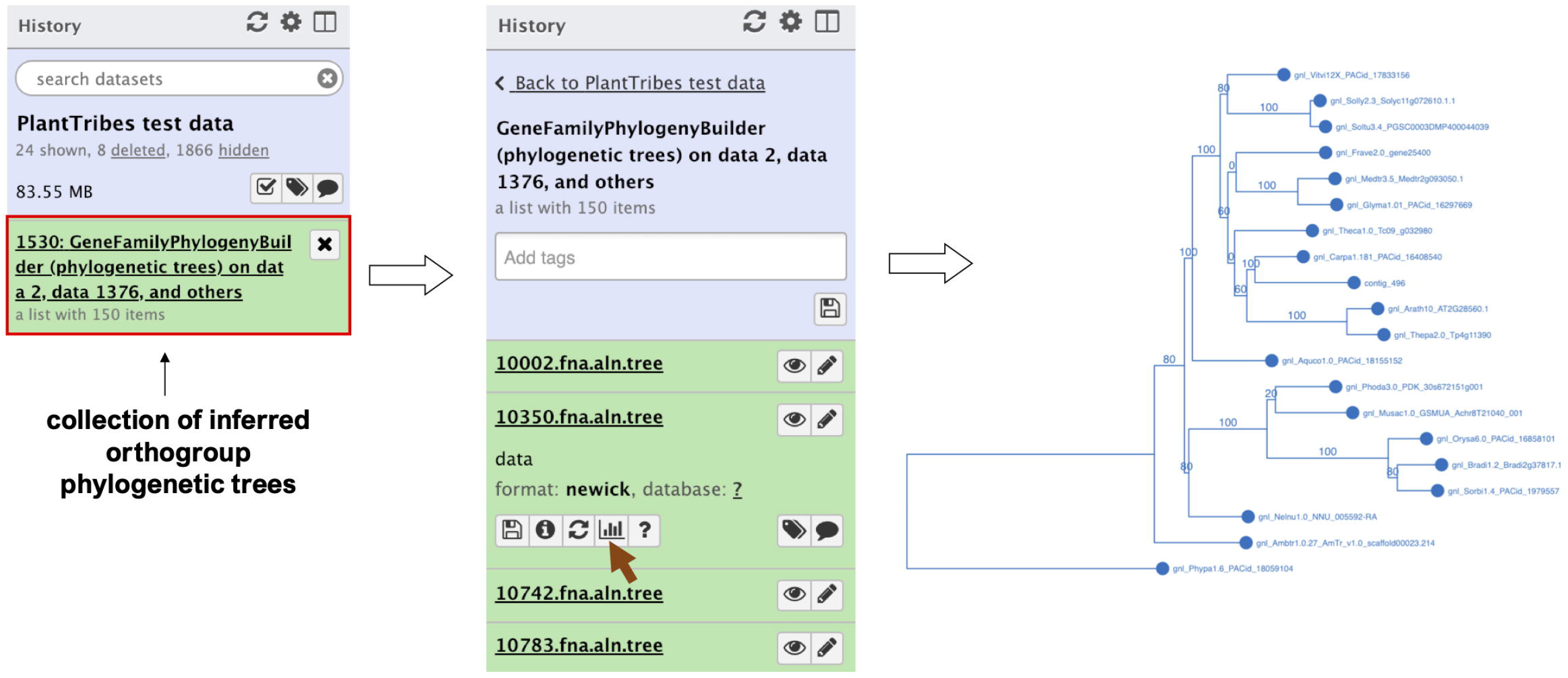
An illustration of an orthogroup phylogenetic tree produced by the Galaxy PlantTribes2 GeneFamilyPhylogenyBuilder using the test dataset. Results can be visualized in Galaxy using either the Phylocanvas (demonstrated here) or the PHYLOViZ plugin.

#### 2.2.5 Estimation of genome duplications

The *KaKsAnalysis* tool estimates paralogous and orthologous pairwise synonymous (*Ks*) and non-synonymous (*Ka*) substitution rates using PAML (Yang, 2007) for a set of protein coding genes (*i*.*e*., produced by the *AssemblyPostProcessor*), with duplicates inferred from the phylogenomic analysis (using both the *GeneFamilyClassifier* and *GeneFamilyPhylogenyBuilder*) or from an external source. Optionally, the resulting set of estimated *Ks* values can be clustered into components using a mixture of multivariate normal distributions, implemented in the EMMIX (McLachlan and Peel, 1999) software, to identify significant duplication event(s) in a species or a pair of species. The *KsDistribution* tool then plots the *Ks* rates and fits the estimated significant component(s) onto the distribution (Figure 4).

**Figure 4:**
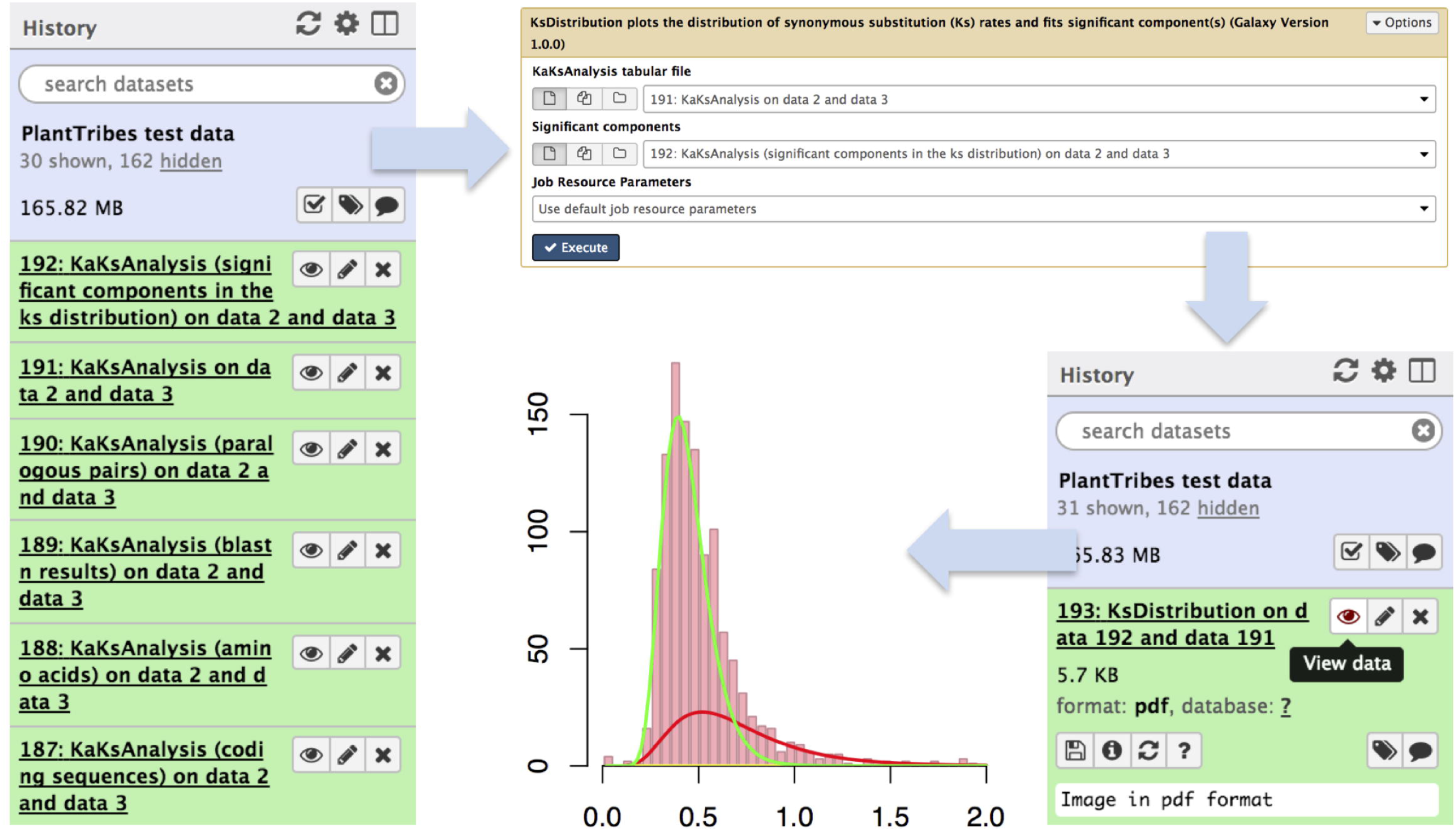
An illustration of genome duplication events detected using the Galaxy PlantTribes2 KaKsAnalysis tool. The KaKs analysis tool produces a list of outputs including self blastn results (item 189), a list of paralogous pairs (item 190), Ka (non-synonymous) and Ks (synonymous) substitution rates (item 191), and the significant components in the Ks distribution (item 192). Then the distribution of estimated paralogous pair Ks values is clustered into components using a mixture of multivariate normal distributions to identify significant duplication event(s) (item 193) and is visualized using Galaxy built-in tools.

## 3. Results

### 3.1 Performance evaluation of sequence classifiers

PlantTribes2 uses BLAST (blastp) and HMMER (hmmscan and hmmsearch) algorithms to classify inferred protein sequences into orthologous gene family clusters, a foundational step for many downstream analyses. To demonstrate the versatility of these two classifiers on gene family clusters, we present evaluations for classification algorithms using the pre-computed 22Gv1.1 gene family scaffold (Supplemental table 2). This scaffold contains annotated protein coding sequences (CDSs) for 22 representative land plant genomes, including nine rosids, three asterids, two basal eudicots, five monocots, one basal angiosperm, one lycophyte, and one moss.

Three taxa with varying evolutionary distances in relationship to all the other taxa in the 22Gv1.1 gene family scaffold were selected: the only moss species, *Physcomitrella patens*, and two asterid sister species, *Solanum lycopersicum* and *Solanum tuberosum*. These three taxa were removed from the scaffold and then classified back to assess recall and precision of the BLAST and HMMER classifiers (Vihinen, 2012). Only protein sequences reassigned to their original orthologous clusters were considered true positives. In addition, F-score, a single metric that considers both recall and precision to measure the overall performance of the two classifiers, was calculated (Vihinen, 2012). The procedure is performed as described below:

**(1) Distant**: *Physcomitrella patens* was removed and sorted back into the scaffold to evaluate the performance of classifiers with distant species. No other moss or bryophyte species are present in this scaffold.

**(2) Moderately Distant**: Both *Solanum lycopersicum* and *Solanum tuberosum* were removed, and *S. lycopersicum* was sorted back into the scaffold to evaluate the performance of classifiers with moderately distant species. After removing both *S. lycopersicum* and *S. tuberosum*, no other sister species in the same plant family are present in the scaffold. However, close lineages, including three asterids and nine rosids, are present in the scaffold.

**(3) Confamilial**: *Solanum lycopersicum* was removed and sorted back into the scaffold to evaluate the performance of classifiers with confamilial species. *Solanum tuberosum*, a sister species from the same plant family, is present in the scaffold.

As shown in Figure 5, the overall classification performance for BLAST and HMMER is similar based on the F-scores across different evolutionary distances (73%-94% for BLAST, 67%-94% for HMMER). In addition, both classifiers have a higher recall rate when classifying into OrthoMCL and OrthoFinder clusters (90% -96%) compared to GFam clusters (58% -80%). HMMER is slightly more sensitive than BLAST when the evolutionary distance is significant, while BLAST is much more sensitive when classifying into GFam clusters at any evolutionary distance. Precision for both classifiers is similar across the evolutionary distance of the scaffold (78% -95%). Classifying into OrthoFinder clusters yields much higher precision (80%-95%) than classifying into OrthoMCL (78%-86%) and GFam (81%-85%) clusters. These findings suggest that, regardless of the sequence classifier algorithm used or evolutionary distance, clusters inferred by orthology methods (OrthoFinder and OrthoMCL) result in better clustering performance compared to clusters inferred by a consensus domain-based method (GFam). We recommend using the merged classification results from BLAST and HMMER, as implemented in the pipeline, because it leverages the strength of both classifiers.

**Figure 5:**
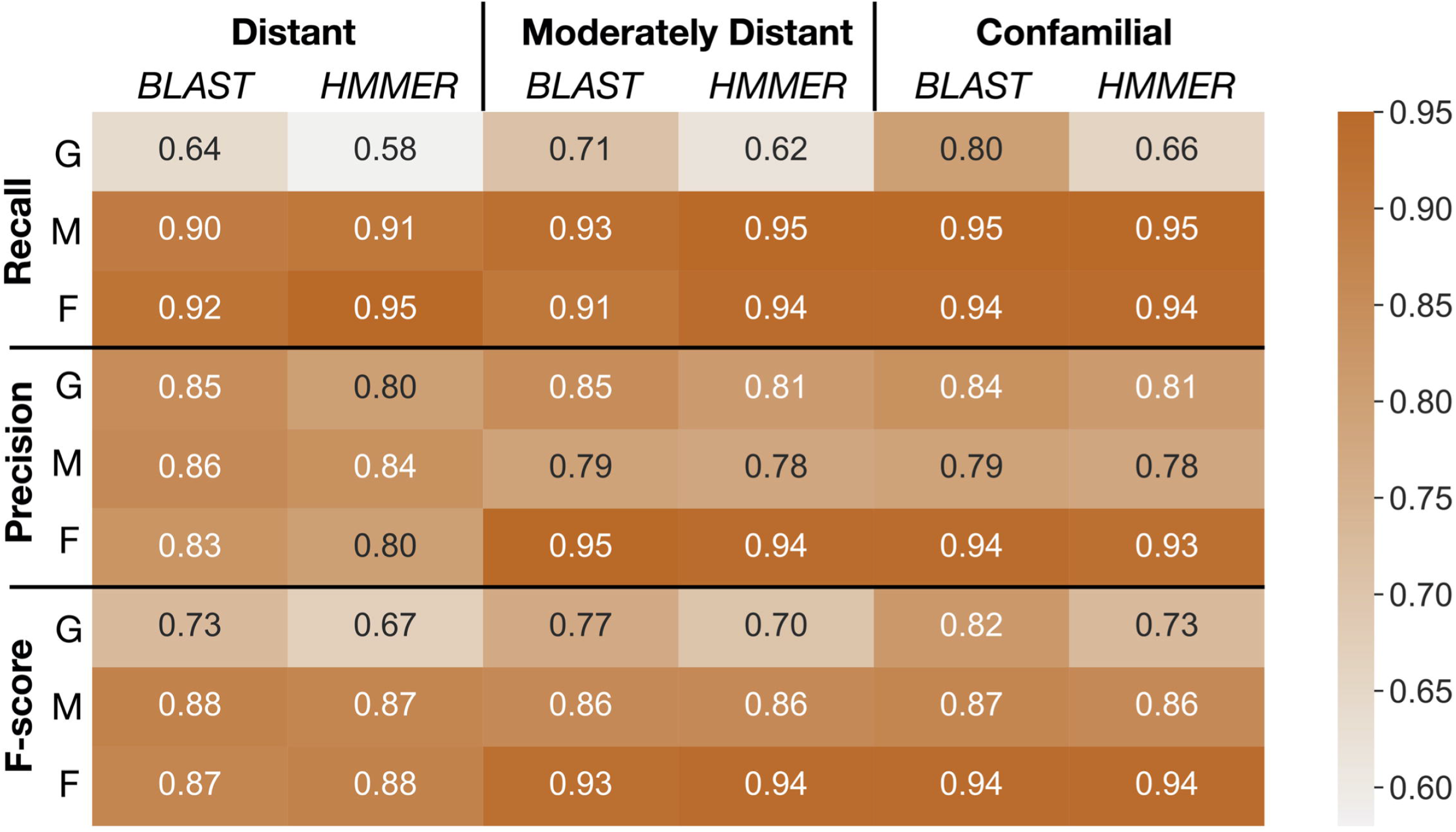
Summaries of performance evaluation of classification rates for BLAST and HMMER classifiers. Recall, precision, and F-score (Vihinen, 2012) for the two classifiers are measured on GFam (G), OrthoMCL (M), and OrthoFinder (F) clustering methods to determine how well taxa at different distances are classified into the PlantTribes2 22Gv1.1 gene family scaffold. Larger values are better. **Distant**: remove and sort back *Physcomitrella patens*, a species distantly related to all other scaffolding species; **Moderately distant**: remove *Solanum lycopersicum* and *S. tuberosum*, then sort back *S. lycopersicum*. No other Solanaceae species are present in the scaffold, but moderately distant species, *i*.*e*., other asterids, are used as scaffolding species; **Confamilial**: *S. lycopersicum* was removed and sorted back. A confamilial species, *S. tuberosum*, is present in the scaffold.

### 3.2 Examples of application

Here we provide examples of how to use PlantTribes2 to answer specific questions regarding (1) alleviating fragmentation issues in a *de novo* transcriptome assembly, (2) evaluation and improvement of gene families and gene models, and (3) assessing the quality of genomes in closely related species.

#### 3.2.1 Evaluation of targeted gene family assembly

*De novo* assembly of RNA-Seq data is commonly used to reconstruct expressed transcripts for non-model species that lack quality reference genomes. However, heterogeneous sequence coverage, sequencing errors, polymorphism, and sequence repeats, among other factors, cause algorithms to generate contigs that are fragmented (Zhang et al., 2014; Honaas et al., 2016). In order to demonstrate the utility of the targeted gene family assembly function in PlantTribes2, we obtained raw Illumina transcriptome datasets sequenced by the Parasitic Plant Genome Project (http://ppgp.huck.psu.edu) that represent key life stages of three parasitic species in the Orobanchaceae family (Westwood et al., 2012; Yang et al., 2015b). These species span the full spectrum of plant parasitism (Westwood et al., 2010, 2012), and include *Triphysaria versicolor, Striga hermonthica*, and *Phelipanche aegyptiaca*. Species-specific transcriptome assemblies were performed with Trinity using two approaches: (1) combining raw Illumina reads from all development stages of the plant in a single assembly, and (2) multiple assemblies of individual developmental stages of the plant. A BUSCO (benchmarked universal single-copy orthologs) (Manni et al., 2021) assembly quality assessment using 1,440 universally conserved land plants’ single-copy orthologs suggests that the assembly combining all raw data recovers more conserved single-copy genes than any developmental stage-specific assembly (Combined *v*.*s*. Stage in Figure 6 and Supplemental table 4). However, a meta-assembly of transcripts from both approaches with the targeted gene family function of the *AssemblyPostProcessor* tool using the 26Gv1.0 gene family scaffold recovers even more full-length conserved single-copy genes (Meta *v*.*s*. others in Figure 6 and Supplemental table 4). Therefore, the meta-assembly implementation of the PlantTribes2 *AssemblyPostProcessor* tool can benefit many comparative transcriptome studies of non-model species to alleviate transcript fragmentation in gene families of interest.

**Figure 6.**
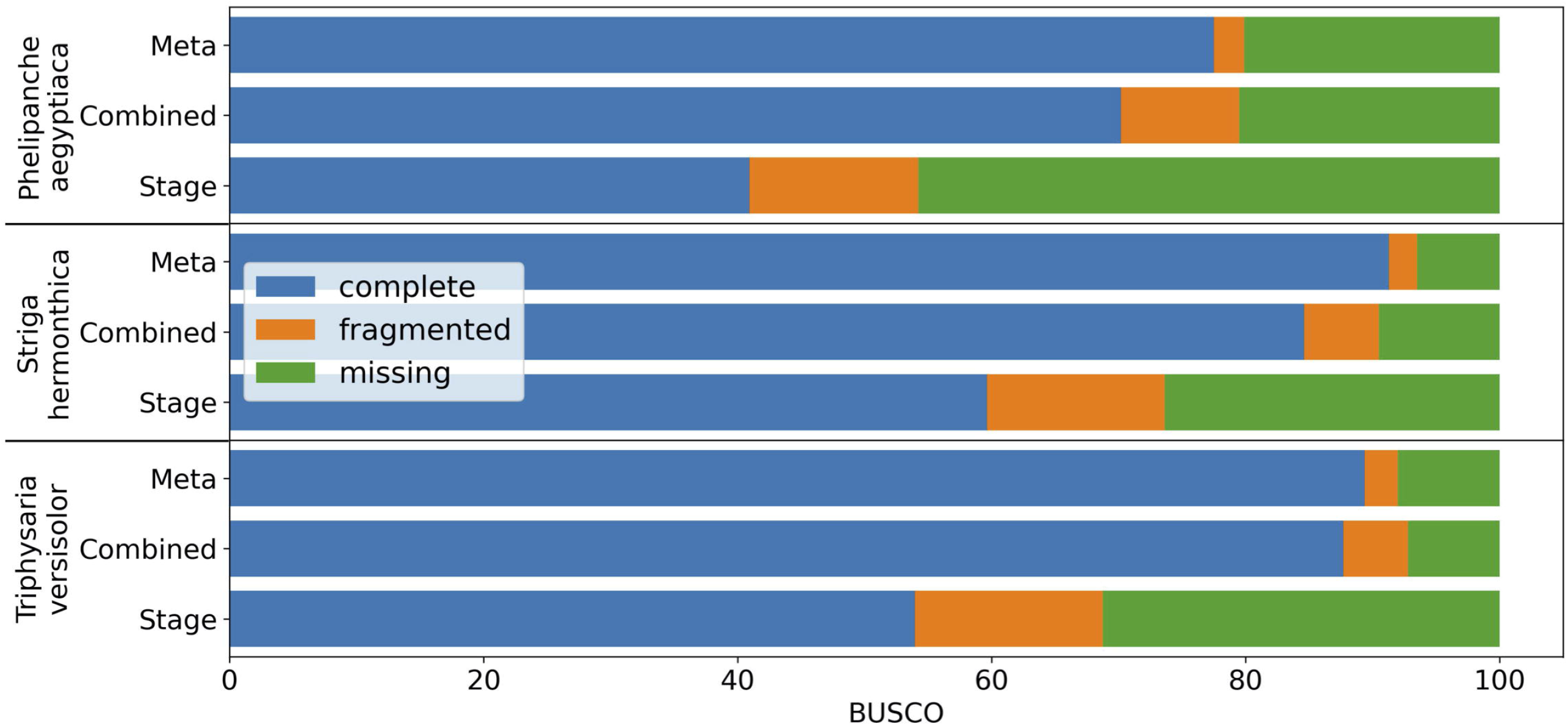
BUSCO completeness assessment of transcriptome assemblies to illustrate the results from targeted gene family assembly (meta-assembly) function in the PlantTribes2 *AssemblyPostProcessor* tool compared to Trinity approaches. Color bars indicate complete (blue), fragmented (orange), and missing (green) BUSCOs. Assemblies of parasitic plants, *Phelipanche, Striga*, and *Triphysaria*, examined include (1) developmental stage-specific assemblies (Stage, only the average of all the stages were shown in the plot), (2) assemblies combining all stage-specific raw data (Combined), and (3) meta-assembly of stage-specific assemblies and combined assembly (Meta) using *AssemblyPostProcessor*.

#### 3.2.2 Application in evaluating and improving gene families

Gene and gene family studies in non-model organisms are challenging due to the varying quality of genome assemblies and annotations, as well as the lack of closely related species as an annotation reference. Thousands of genes lack accurate gene models in draft and early version genomes (Darwish et al., 2015; Marx et al., 2016; Li et al., 2017, 2019; Pilkington et al., 2018; Liu et al., 2021) creating pitfalls for global-scale analyses, but especially for researchers conducting reverse genetics studies. For example, in the first version of the apple (*Malus domestica*) genome annotation, we discovered that the gene model of *MDP0000250518*, annotated as *MdPIN8a* by Song *et al*., (2016), is problematic. A nucleotide sequence comparison of *MDP0000250518* and its orthologous genes in other Rosaceae genomes, identified using the PlantTribes2 orthogroup classification function, showed that this gene model is likely a combination of the putative *MdPIN8a* and a neighboring gene, which encodes a voltage dependent anion channel (VDAC) (Figure 7). These two genes are located about 3000bp apart on the same chromosome in most Rosaceae genomes (Supplemental table 5). Analyses carried out using this incorrect gene model may confound or compromise the work. For example, in absence of the contextual gene family information we now have from analyses with PlantTribes2, the authors in Song et al., (2016) unknowingly designed primers for the *MDP0000250518* gene model that targeted the *VDAC* gene rather than the actual gene of interest, *MdPINB8a* (Figure 7). We identified the mis-annotated gene using contextual gene family information ; a reliable way to avoid such pitfalls.

**Figure 7.**
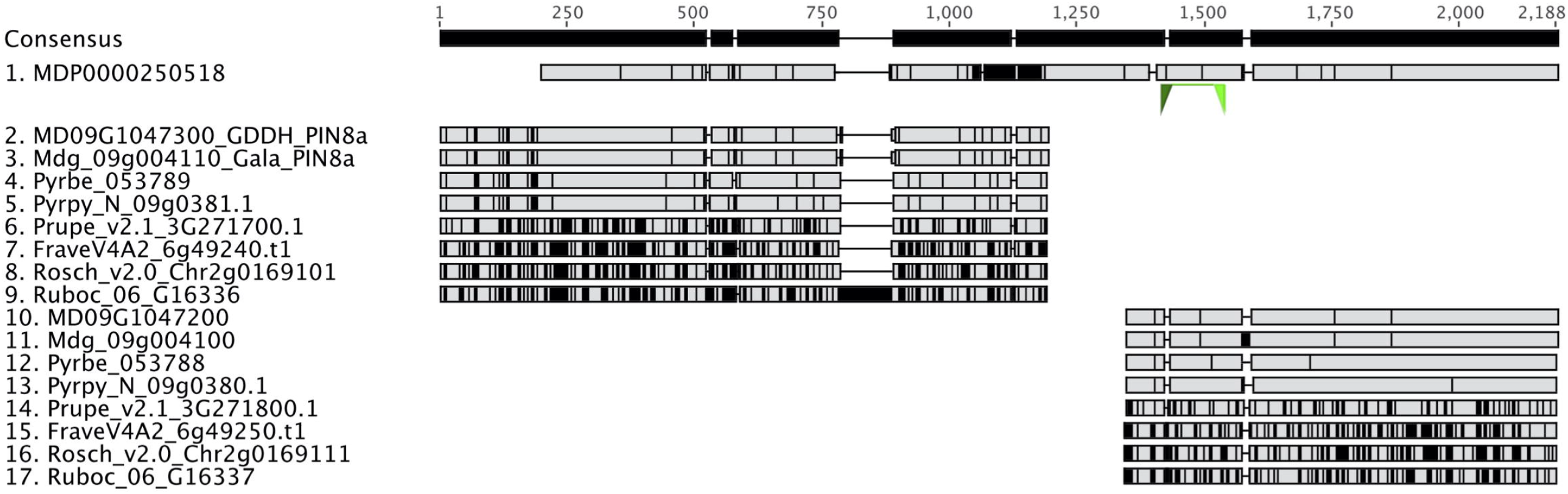
Identification of an incorrect auxin transporter gene model, *MdPIN8a*, in *Malus domestica* genome annotation version 1. Nucleotide sequence alignment of putative *PIN8a* and *VDAC* genes from *Malus* and *Pyrus. MDP0000250518* (sequence 1) gene model is a combination of two genes. The 5’ end of *MDP0000250518* shares high sequence similarity with the *PIN8a* gene from other Rosaceae species (sequence 2 to 9), while its 3’ end shows evidence of homology to a neighboring gene, *VDAC*, in the investigated genomes (sequence 10 to 17). Green triangles below *MDP0000250518* show the binding sites of the qRT-PCR primers used in the Song *et al*. 2016 research. Gray color indicates identical nucleotides compared to the consensus, while black color indicates different nucleotides.

Better gene models can be obtained from re-annotating existing or new genome assemblies with additional transcriptome data. For instance, tens of thousands of gene models were improved or added in the subsequent annotations in several strawberry genomes (Darwish et al., 2015; Li et al., 2017, 2019; Liu et al., 2021). In later versions of apple genome annotations, erroneous gene models such as *MdPIN8a* and the neighboring gene, *VDAC*, are corrected and are now concordant with other Rosaceae (Figure 7). This improved gene information provides a better starting point for studies like Song et al. 2016, however, full reannotation of complex plant genomes is a time-consuming and a resource-intensive undertaking.

A more efficient solution is targeted gene model improvement by evaluation of genes of interest (GOIs) from a gene family perspective. The comparative genomic and phylogenomic tools offered by PlantTribes2 allows researchers to efficiently compare orthologous genes across many closely related species and identify problematic genes in a high-throughput fashion. In a recent study with a goal to identify tree architecture genes in *Pyrus* (pear), functions from PlantTribes2 were used at the core of the workflow (Zhang et al., 2022, the companion paper in this issue). Using the alignments and phylogenies generated by the *GeneFamilyAligner* and *GeneFamilyPhylogenyBuilder* tools from PlantTribes2, hundreds of problematic gene models were identified. For instance, two fragments of a putative pear *DWARF4* (*DWF4*) gene were found in the *Pyrus communis* ‘Bartlett’ Double Haploid (Bartlett.DH) genome annotation (Linsmith et al., 2019), one of which showed little evidence of homology at the 3’ end of its coding sequence compared to other apple and pear *DWF4* genes. This problem was easily recognized in the nucleotide sequence alignment and phylogeny produced by PlantTribes2 (Figure 8A and C). Moreover, the homologous sequences from the PlantTribes2 orthogroups were readily used as resources for target-gene family annotation tools, such as TGFam-Finder (Kim et al., 2020) and Bitacora (Vizueta et al., 2020). In the case of *DWF4*, using the PlantTribes2 derived orthogroup information as reference, a more complete *DWF4* gene homologous to other Maleae sequences was annotated from the Bartlett.DH genome (Figure 8 B and C). More examples like the *DWF4* gene are presented in Zhang et al., 2022.

**Figure 8.**
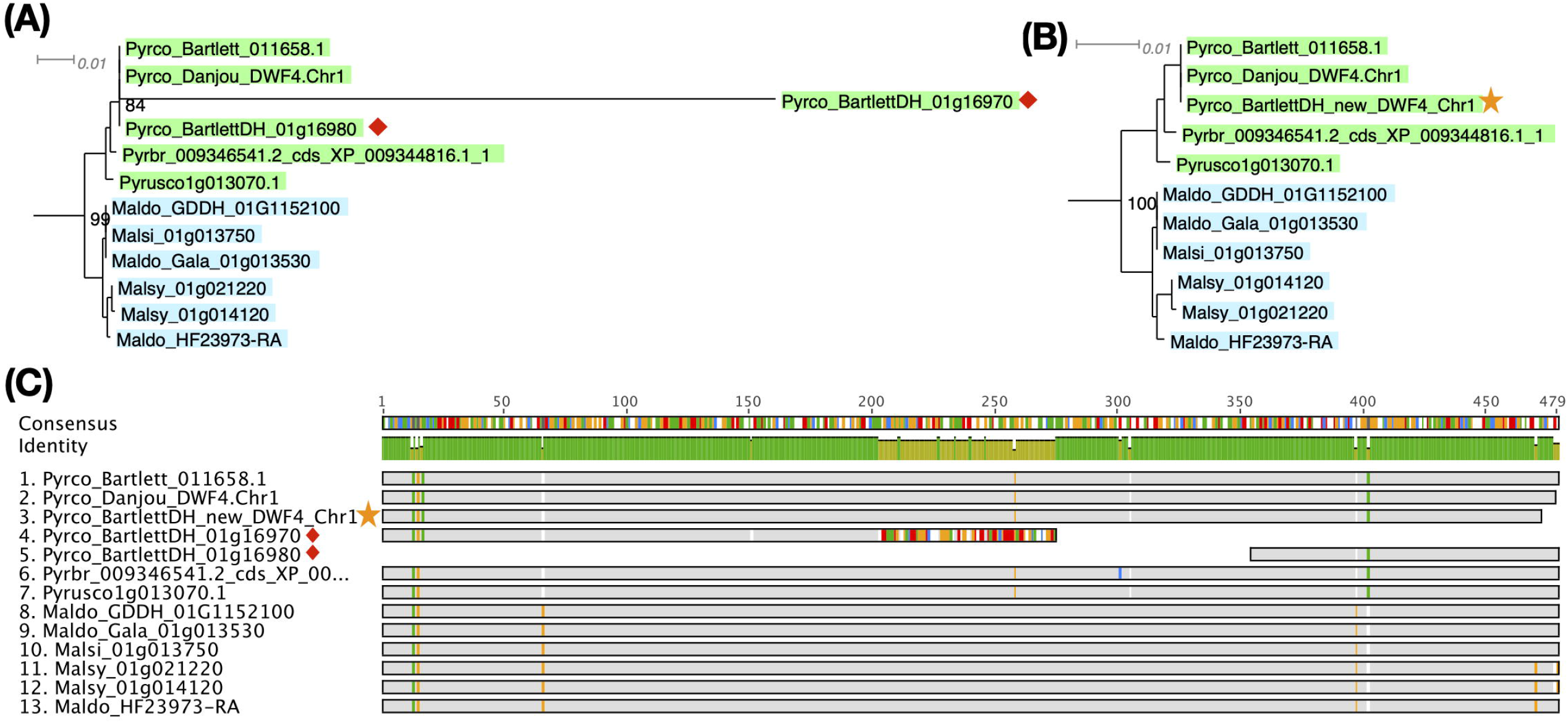
Example of a putative *DWF4* gene before (red diamond) and after (yellow star) improvement. **(A)** and **(B)** show a section of the *DWF4* gene family tree with Bartlett_DH gene models before and after improvement, respectively. **(C)** is the sequence alignment of the *DWF4* gene coding region in *Malus* and *Pyrus* genomes. Gray color indicates identical nucleotides compared to the consensus, while colors indicate different nucleotides.

#### 3.2.3 Application in evaluating genome quality

A BUSCO analysis is a widely accepted benchmark to assess the completeness and accuracy of genomic resources (Manni et al., 2021). However, it only takes into consideration a very small fraction of the gene space. By definition, BUSCOs appear as highly conserved single copy genes in many organisms and return rapidly to single copy following gene and genome duplication. BUSCO genes may not reflect the quality of more challenging regions of the genome and the integrity of complex and divergent gene families. With more genomic resources being produced, especially in some agronomically important genera/species, lineage-specific BUSCO databases have been developed, bringing in larger numbers of markers. For instance, the poales_odb10 contains 3 times more markers than the generic embryophyta_odb10. However, this type of database has only been developed for 4 plant orders (Brassicales, Solanales, Poales, and Fabales), and like other BUSCO databases, only single copy genes are used. Following the same philosophy as the lineage-specific BUSCO databases, the natural next step is a gene-by-gene assessment on a genome scale, as proposed by Honaas *et. al*., (2016) regarding *de novo* transcriptome assembly evaluation. Here we present a case study of using the objective orthogroup classification offered by PlantTribes2 to evaluate the quality of genome annotations from a comparative perspective in Rosaceae, a step towards a gene-by-gene approach.

The number of publicly available Rosaceae genomes, generated by researchers all around the world using different technologies, has increased exponentially in the last decade (Jung et al., 2018). To better estimate the accuracy and sensitivity of genome annotation across a wide range of Rosaceae species, we created family-specific “CoRe OrthoGroups (CROGs) -Rosaceae”. First, 26 representative genomes from six genera (*Malus, Pyrus, Prunus, Fragaria, Rosa*, and *Rubus*. Supplemental tables 6 and 7) in five major Rosaceae tribes were classified into the PlantTribes2 26Gv2.0 scaffold. Next, the union of orthogroups from each genus was generated, creating genus-level master orthogroups. Then the overlap of the six master orthogroups, consisting of 9656 orthogroups, were designated as the CROGs (Figure 9A, Supplemental table 8), which is so far the most complete list of core Rosaceae genes. Rich information from the CROGs, *i*.*e*., the percentage of CROGs captured in each genome, gene counts in CROGs, and sequence similarity compared to the CROG consensus, can be used to assess annotation quality, pinpoint areas needing improvement, and find potentially interesting biology.

**Figure 9.**
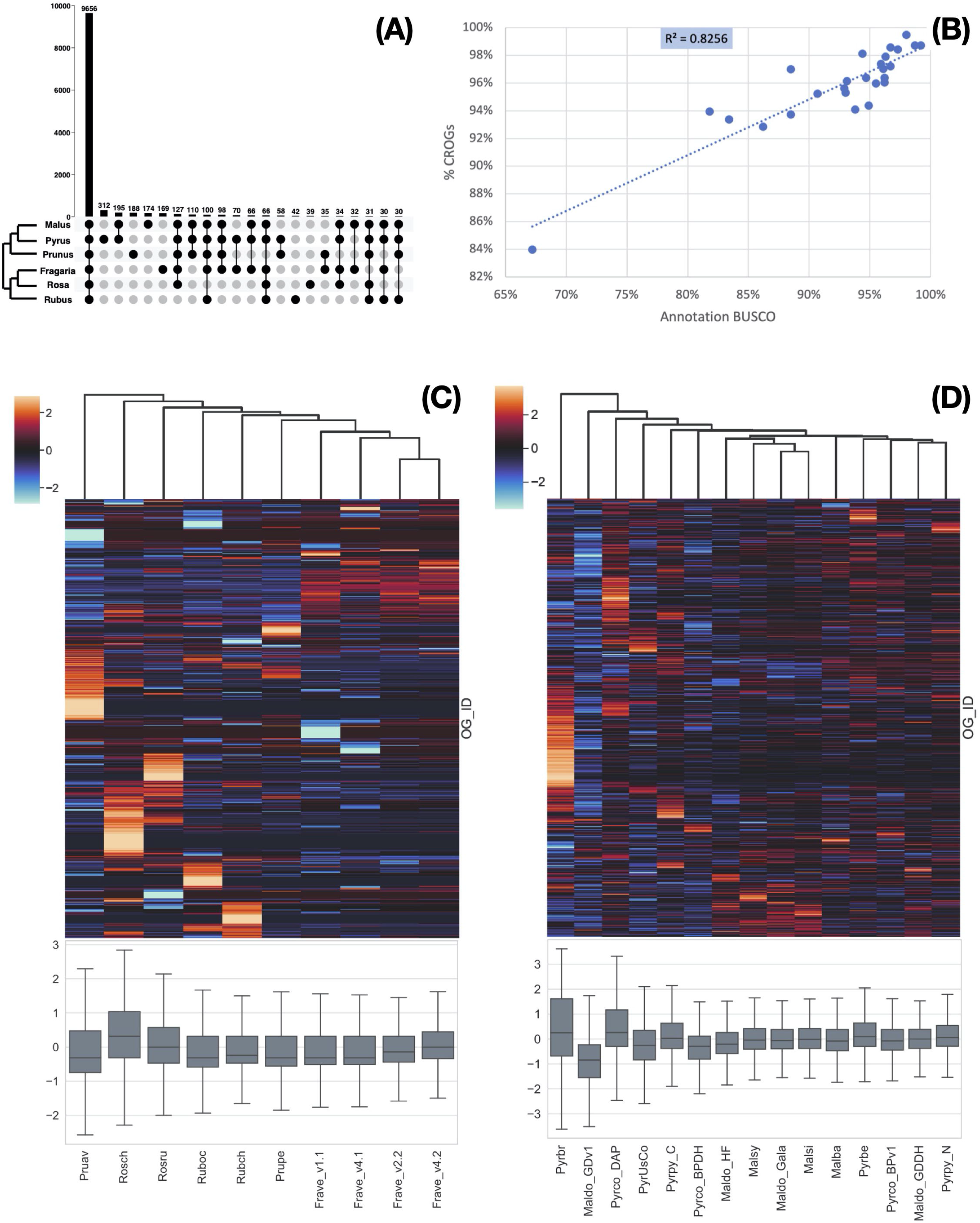
CoRe OrthoGroups - Rosaceae (CROGs). **(A)** Upset plot showing overlapping orthogroups between six Rosaceae genera, including 9656 orthogroups shared by all six genera (designated as “CROGs – Rosaceae”). **(B)** High correlation between Rosaceae genome annotation BUSCOs and % CROGs captured in the genomes (p<0.01). **(C)** Z-score distribution of gene counts in CROGs among selected Rosaceae genomes excluding *Malus* and *Pyrus*, shown as a clustermap (upper) and a box plot (lower). Each column represents a genome and each row in the clustermap represents a CROG. **(D)** Z-score distribution of gene counts in CROGs among selected *Malus* and *Pyrus* genomes, shown as a clustermap (upper) and a box plot (lower). Genome abbreviations can be found in Supplemental table 7.

First, we calculated the percentage of CROGs captured in 26 Rosaceae genomes and correlated the %CROGs with the corresponding annotation BUSCO scores (Supplemental table 6). The high positive correlation (R^2^ = 0.82, Figure 9B) indicates that these two philosophically similar approaches draw the same conclusions for most genomes, however, CROGs provide additional information allowing more in-depth explorations of annotation quality.

Next, we calculated gene counts in each CROG. Due to the difference in chromosome numbers (17 chromosomes in Maleae and 9 in other genera) and a unique recent whole genome duplication event in the common ancestor of the Maleae (Hodel et. al., 2022), apple and pear genomes have more gene copies in most orthogroups than other Rosaceae. To make more appropriate comparisons, we generated two CROG gene count matrices, one for Maleae and one for other Rosaceae (Supplemental tables 9 and 10, respectively). Our hypothesis is that a high-quality genome will have a predictable and consistent number of genes in a large majority of CROGs. This is because issues that have predictable impacts on genome assembly and annotation are dependent on individual genome characteristics, the data used in assembly and annotation, and the various methodologies employed therein -thus creating a comparative framework with complementary error structure. Simply put, it is unlikely that a gene family will show a consistent yet erroneous shift in gene content due to methodological reasons alone. This perspective can reduce the false positive rate for evolutionary inference of lineage-specific shifts in gene family content by flagging changes in individual genomes that may be due to methodological bias.

As expected, in the non-Maleae matrix, nearly half of the CROGs (4,728) have the same number of genes or different gene counts in only 1 or 2 genomes. When we visualized the gene count matrix using the Seaborn z-score clustermap package (CROGs with standard deviation of 0 were removed prior to plotting), the four different versions of *Fragaria vesca* annotations clustered together (Figure 9C). They shared similar z-score patterns in most CROGs, but fewer low z-score regions (shown as cooler colors) were found in the later versions of annotation (v2.2 and v4.2). These two annotations also have a mean z-score closer to 0 and relatively small variance compared to the earlier annotations. A similar pattern was seen while comparing the first version of the apple genome, Maldo_GDv1, to the more recent ones (Maldo_Gala and Maldo_GDDH in Figure 9D). Our results are consistent with previous reports (Daccord et al., 2017; Li et al., 2017, 2019) and the CROG approach provided a fast and easy-to-visualize way to summarize these findings.

The clustermaps also allowed us to gain new insights from these genomes. For instance, there is not clear clustering of *Malus* or *Pyrus* at the genus level, however, the more recent genomes, which have less variable z-score distribution centered near 0, are clustered together (Figure 9D). We hypothesize that the current clustering is mainly driven by genome annotation strategy and quality, and therefore it is showing methodological similarities rather than biological patterns. The fact that *Malus domestica* Gala (Maldo_Gala) is clustered with *M. sieversii* (Malsi) and *M. sylvestris* (Malsy), genomes generated using the same method, rather than the other high-quality *M. domestica* genome, Maldo_GDDH, supports this hypothesis (Daccord et al., 2017; Sun et al., 2020).

Another unexpected observation is that the earlier version of the European pear (*Pyrus communis*) genome, Pyrco_BPv1 (Chagné et al., 2014), shared a more similar gene count pattern with some of the best Maleae genomes. On the contrary, the second version, Pyrco_BPDH (Linsmith et al., 2019), a double haploid genome, does not. Apple and pear are highly heterozygous, which is known to cause fragmented genome assembly and introduce multiple alleles to the annotation. Sequencing isogenic genotypes, such as a double haploid, is a common solution (Linsmith et al., 2019; Daccord et al., 2017; Zhang et al., 2019). This process will reduce the complexity in genome assembly and should have little to no influence on the number of genes in a genome, or even in individual gene families. When a smaller number of protein coding genes were annotated from the Pyrco_BPDH genome compared to version one, the authors hypothesized that the difference is resulted from removal of allelic sequences annotated as genes (“allelic genes”) in the much more contiguous double haploid genome (Linsmith et al., 2019). However, our CROG gene count matrix indicates that the smaller gene number in Pyrco_BPDH is caused, at least in part, by CROG genes and gene families missing from the annotation -indeed the Pyrco_BPv1 genome captured a vast majority of the CROGs with the expected gene count, despite annotation of some allelic genes. This statement is supported by an investigation in putative tree architecture gene families by Zhang *et al*., 2022 (the companion paper). About half of the genes of interest were missing in the original Pyrco_BPDH annotation, but were recovered using a polished assembly and targeted annotation approaches (Figure 10).

**Figure 10.**
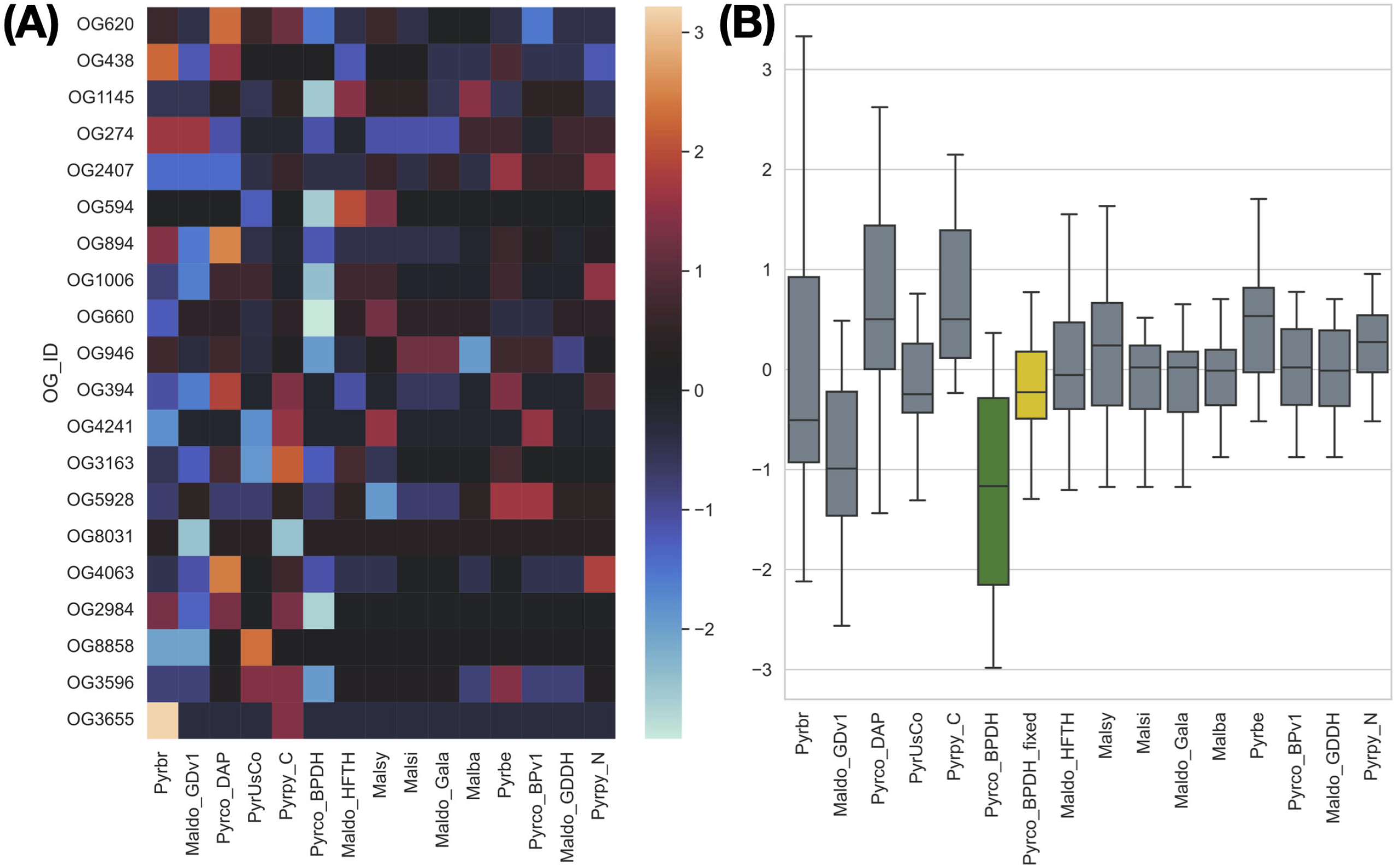
The gene count z-score of selected tree architecture gene families across *Pyrus* and *Malus* genomes. Pyrco_BPDH orthogroups have lower z-scores than most others, which is shown with a cooler color in the heatmap **(A)** and lower average z-score (green box in **B**), indicating fewer than expected gene counts. These missing genes were discovered after the targeted re-annotation process, which brought the average gene count z-score closer to 0 (yellow box in **B**) and comparable to other high-quality genomes.

The “hot” zones in the clustermaps also attract attention. To investigate the hot zones in the Maleae matrix, we examined the gene counts and annotation of 150 CROGs with the highest z-score from each genome. In most genomes, these CROG annotations lack a pattern, and the high z-score is caused by one or few extra copies, which may be caused by the introduction of alleles from fragmented assembly or could indicate genome-specific duplications. However, in some high z-score CROGs in *Pyrus betulifolia* (Pyrbe) (Dong et al., 2020), *Malus sieversii* (Msi), *M. sylvestris* (Msy), and *M. domestica* ‘Gala’ (Maldo_Gala) (Sun et al., 2020), the targeted genome has up to 10 times more genes than the others and the annotation of these CROGs are often related to transposons and repeat-containing genes (Supplemental table 11). This finding suggests certain downstream analyses, such as repeat type comparison and gene family expansion estimation, can be bolstered against such pitfalls by a CROG analysis.

Using the PlantTribes2 orthogroup classification, we created a new method to evaluate genome quality in more depth, leveraging resources across an important plant family. The CROG gene count matrix does not only provide a highly effective way to visualize differences in gene numbers from a comparative genomic perspective, but also pinpoints where improvements could be made. As genomic resources are rapidly increasing, a CROG analysis can also help to inform the selection of the most appropriate genomes for comparative genomic studies, by avoiding specific issues related to assembly and annotation. Moreover, this approach can be applied to any groups of plants, creating custom CROGs for assessing the quality of genomes of interest.

## 4. Conclusions

PlantTribes2 uses pre-computed or expert gene family classifications for comparative and evolutionary analyses of gene families and transcriptomes for all types of organisms. The two main goals of PlantTribes2 are: (1) continual development of a scalable and modular set of analysis tools and methods that leverage gene family classifications for comparative genomics and phylogenomics to gain novel insight into the evolutionary history of genomes, gene families, and the tree of life; (2) to make these tools broadly available to the research community as a stand-alone package and also within the Galaxy Workbench. Many genomic studies, including inference of species relationships, the timing of gene duplication and polyploidy, reconstruction of ancestral gene content, the timing of new gene function evolution, detection of reticulate evolutionary events such as horizontal gene transfer, assessment of gene family and genome quality, and many others, can all be performed using PlantTribes2 tools. The modular structure, which allows component tools of the pipeline to be independent from each other, makes the PlantTribes2 tools easy to enhance over time.

## 5. Data Availability

Project name: PlantTribes2 Archived version: 1.0.4

Project home page: https://github.com/dePamphilis/PlantTribes

Galaxy: https://usegalaxy.org

Bioconda: https://bioconda.github.io/search.html?q=PlantTribes

Tutorials: https://github.com/dePamphilis/PlantTribes/blob/master/docs/Tutorial.md. https://galaxyproject.org/tutorials/pt_gfam/

Operating system(s): Linux, Mac OS X Programming language: Perl, Python

Other requirements: Web browser for Galaxy License: GNU

## 6. Acknowledgements

We acknowledge funding from the National Science Foundation (NSF) DBI-1238057, NSF IOS-0922742, the NSF Plant Cyberinfrastructure Program through iPlant (now CyVerse; DBI-

0735191), the The iPlant Tree of Life Grand Challenge Project, USDA ARS, WTFRC grant AP-19-103. The authors thank Heidi Hargarten for editing and revising the manuscript.

## 8. Author contributions

E.K.W., J.L.-M., and C.W.D conceived and designed the research. E.K.W, H.Z, L.A.H, and G.VK performed the analyses. All authors participated in writing and revising the manuscript.

## 9. Contribution to the field statement

The field of functional genomics aims to link genes to traits, but at a massive scale. However, most plants are hard to study, such as fruit trees -they are large and can take many years before producing fruits and seeds, making molecular biology experiments in these plants that aim to directly examine gene function very difficult. Instead, researchers tend to study the functions of genes in plants that are amenable to laboratory experiments, like rice, tomato, and a small cousin of the mustard plant - Arabidopsis. This creates a need to transfer knowledge from well-studied plants to important agricultural crops. A method to do this starts by sorting genes into a family tree, which helps us learn about the genes in crops that have distant cousins in, for instance, tomato. Importantly, it allows us to transfer experimental knowledge between plants. This in turn accelerates studies aimed at learning about the genes that control traits in important crops, like fruit trees, that are difficult to study in the lab. The software pipeline we describe in our manuscript, PlantTribes2, is a user-friendly way to sort genes into gene families so we can more easily transfer gene knowledge between plants.

